# Passive diffusion through nuclear pore complexes regulates levels of the yeast SAGA and SLIK coactivators complexes

**DOI:** 10.1101/710020

**Authors:** Tadashi Makio, Richard W. Wozniak

## Abstract

Nuclear pore complexes (NPCs) control gene expression by regulating the bi-directional exchange of proteins and RNAs between nuclear and cytoplasmic compartments, including access of transcriptional regulators to the nucleoplasm. Here we show that the yeast nucleoporin Nup170, in addition to binding and silencing subtelomeric genes, supports transcription of genes regulated by the SAGA transcriptional activator. Specifically, we show that less SAGA complex is bound to target genes in the absence Nup170. Consistent with this observation, levels of the SAGA complex are decreased in cells lacking Nup170, while SAGA-related SLIK complexes are increased. This change in the ratio of SAGA to SLIK complexes is due to increased nuclear activity of Pep4, a protease responsible for production of the SLIK complex. Further analyses of various nucleoporin mutants revealed that the increased nuclear entry of Pep4 observed in the *nup170*Δ mutant likely occurs as consequence of an increase in the sieving limits of the NPC diffusion channel. On the basis of these results, we propose that changes in passive diffusion rates represents a mechanism for regulating SAGA/SLIK complex-mediated transcriptional events.

## Introduction

The nuclear envelope double membrane forms an impermeable barrier between the nucleoplasm and the cytoplasm. For molecules to enter or leave the nucleus they are funnelled through massive channels formed by nuclear pore complexes (NPC)s that perforate the nuclear envelope at numerous positions along the surface of the nucleus. NPCs are present in all eukaryotes and their overall structural organization appears largely conserved. In the yeast S. cerevisiae, NPCs are ∼100 nm diameter and consist of multiple copies of ∼30 different proteins, termed nucleoporins or Nups (Aitchison and Rout, 2012). The components are categorized into several classes based on their sequence and functional features. Many of the most conserved Nups form the cylindrical scaffold of the NPC, while a second group, each rich in the phenylalanine-glycine repeat-containing peptides (FG-Nups), line the transport channel and extend into the cytoplasm and nucleoplasm (Aitchison and Rout, 2012).

There are two mechanisms by which molecules move through the NPC: facilitated and passive diffusion. Facilitated diffusion is mediated by nuclear transport factors (NTFs). NTFs contain multiple binding interfaces that allow them to bind nuclear import or export signals in transport cargos as well as the FG-Nups. Interactions of the NTFs with the FG-Nups are generally of low affinity, allowing NTFs and their attached cargos to partition into and out of the NPC channel without an appreciable change in free energy (Rout et al., 2003; Wente and Rout, 2010). Once in the channel, the NTF-cargo complex can diffuse into either cytoplasm or nucleoplasm (Lowe et al., 2010). This process allows cargos of a broad range of sizes to pass through NPCs. Directionality of transport (i.e. import or export) it then established by additional factors (e.g. RanGTP) that bind and dissociate NTF-cargo complexes in the target compartment (Aitchison and Rout, 2012).

Macromolecules lacking nuclear transport signals are also able to enter the nucleus by passive diffusion, but their movement through NPCs is limited by their size. Classical studies using inert dextrans and labeled proteins concluded that NPCs contained a passive channel of 45-59 Å in radius that would allow molecules up to ∼40-60 kD in mass to diffuse through NPCs (Peters, 1983; Paine, 1975; Paine and Feldherr, 1972). Numerous follow up studies supported these conclusions. However, other studies suggested the passive channel could accommodate larger molecules (Wang and Brattain, 2007; Popken et al., 2015). These observations called into question previous assumptions that the passive diffusion channel was rigid, with a defined size threshold. Importantly a recent study showed that the passive diffusion channel is unlikely to be rigid (Timney et al., 2016). Instead these authors proposed that the channel is flexible and capable of allowing passive diffusion of much larger molecules than previously assumed, at rates that decrease with increasing mass. Consistent with this conclusion, analysis of the vertebrate nuclear and cytoplasmic proteome revealed many proteins and protein complexes of up to ∼ 100 kD are equally distributed in both compartments suggesting they partition across the NE (Wühr et al., 2015). The molecular characteristics of the channel that establish and regulate passive diffusion properties are currently unclear. However, what is known is that several FG-Nups present within the central channel of the NPC (Hülsmann et al., 2012; Popken et al., 2015; Timney et al., 2016) and Nups present within the inner ring complex, namely Nup170 and Nup188 (Shulga et al., 2000), are required for maintaining the permeability properties of the passive diffusion channel.

By regulating macromolecular transport, NPCs directly influence the contains of the nucleoplasm and essentially all nuclear processes. Principle among these is transcription of mRNA. Many components of the transcriptional machinery contain NLSs and they have been shown, or assumed, to be imported into the nucleus by NTFs. However, passive diffusion is also likely to contribute to transcriptional control. For example, increases in the mass threshold for passive diffusion through NPCs have been reported to occur in response to stress or as cells age (D’Angelo et al., 2009; Mason et al., 2005), and these changes are accompanied by alterations in transcription (López-Otín et al., 2013). A hypothesis these arises from these observations is that changes in passive diffusion offer a potential mechanism for globally regulating transcription and potentially other nuclear functions. However, what pathways and specific molecular events might be altered in response to changes in passive diffusion remains to be determined.

The impact of Nups on gene expression extends beyond their roles on regulating transport. Various nups have been shown to interact with chromatin and influence the transcriptional status resident genes. In certain cases Nups, including several FG-Nups, have been shown to interact with and assist in the assembly of transcription machinery on actively transcribed genes (Ptak et al., 2014; Ptak and Wozniak, 2016; Ibarra and Hetzer, 2015). In other instances, Nups can play a repressive role (Therizols et al., 2006; Van De Vosse et al., 2013). For example, yeast Nup170 interacts with subtelomeric chromatin, and this association is required for the assembly of silencing factors on chromatin and the repression of resident subtelomeric genes (Van De Vosse et al., 2013). As these varied functions of Nups would imply, the consequences of loss-of-function mutations of a given Nup on gene expression are a product of both direct effects on chromatin structure and as well as indirect effects arising from alteration of the nuclear transport pathway discussed above. Thus, it is important to consider that transcriptional changes observed in a given *nup* mutation are an amalgamation of changes caused by the loss of multiple functions of that Nup. For example, our previous analysis cells lacking Nup170 showed that the majority of genes effected showed an increase in expression, and many of these were detected in regions of the genome that bound Nup170 (Van De Vosse et al., 2013). However, a smaller group of genes showed a decrease in expression suggesting an alternative function for Nup170 in regulating their expression.

Here we have investigated the mechanism by which Nup170 supports the transcription of a group of genes controlled by Spt-Ada-Gcn5 acetyltransferase (SAGA) coactivator complex. We observed that representative genes within this group of show reduced levels of associated SAGA complex in the absence of Nup170. Consistent with these data, cellular levels of SAGA complex were also lower, while a related complex, the SLIK (<u>S</u>AGA-<u>lik</u>e) complex was increased. We show that this change in the ratio of SAGA to SLIK complexes in the *nup170*Δ mutant is due to increased nuclear activity of Pep4, a protease responsible for production of the SLIK complex. Our further analyses lead us to propose that the increased nuclear levels of Pep4 observed in the *nup170*Δ mutant occurs as consequence of increased passive diffusion through NPCs. On the basis of our observations, we propose that changes in passive diffusion rates through NPCs, which can arise as a consequence of changes in cell physiology, represents a mechanism for globally modulating nuclear functions, including the regulation of SAGA/SLIK complex-mediated transcriptional events.

## Results

### Expression of SAGA complex target genes is decreased in *nup170Δ* cells

In a previous study, we found that the gene expression pattern of yeast cells lacking the NPC protein Nup170 showed specific alterations (Van De Vosse et al., 2013). Notably many genes within subtelomeric chromatin regions were upregulated. However, a subset of genes positioned throughout the genome exhibited reduced mRNA levels. Further inspection of these genes revealed an enrichment of genes regulated by SAGA transcriptional coactivator complex (Fig. S1). The SAGA complex contains multiple functional groups, including a histone acetyltransferase, that function to support the transcription of ∼10% of the genes transcribed by RNA polymerase II (Pol II) (Huisinga and Pugh, 2004; see Fig. S1). Genes in this category, referred to as SAGA-dominated genes, are overrepresented among those downregulated in the *nup170Δ* mutant, where 38 genes out of top 100 most downregulated genes are categorized as SAGA-dominated genes (*p* = ∼ 10^−25^ on the Pearson’s chi-squared test).

Levels of mRNAs encoded by SAGA-dominated genes in the *nup170Δ* mutant were further confirmed by qPCR. Analysis of selected SAGA-dominated genes, including *PHO5, PDR5*, and *HIS4*, showed reduced mRNA levels in the *nup170Δ* mutant, while expression levels of several TFIID-dominated genes, including such genes as *DPS1* and *SUR4*, showed no significant change relative to the WT controls (Fig. 1A). Consistent with changes in mRNA levels of SAGA-dominated genes, we also observed a reduction in RNA Pol II density on these genes in the *nup170Δ* mutant. A ChIP experiment using antibodies against the RNA Pol II component Rpb3 was performed to examine RNA Pol II density along two SAGA-dominated gene, *HIS4* and *PDR5*, showing reduced transcripts in the *nup170Δ* mutant. As shown in Fig. 1B, the enrichment of RNA Pol II at the *HIS4* and *PDR5* gene loci was significantly reduced in the *nup170Δ* mutant, suggesting RNA Pol II association with these SAGA-dominated genes was defective. By contrast, three TFIID-dominated genes, *ACT1* and *RPL3*, whose transcript levels were not significantly changed in the *nup170Δ* mutant, show similar levels of RNA Pol II enrichment in the WT and *nup170Δ* mutant strains (Fig. 1B). These results suggest that transcription of the *HIS4* and *PDR5* genes, and likely additional SAGA-dominated genes, is compromised in cells lacking Nup170.

**Figure 1.**
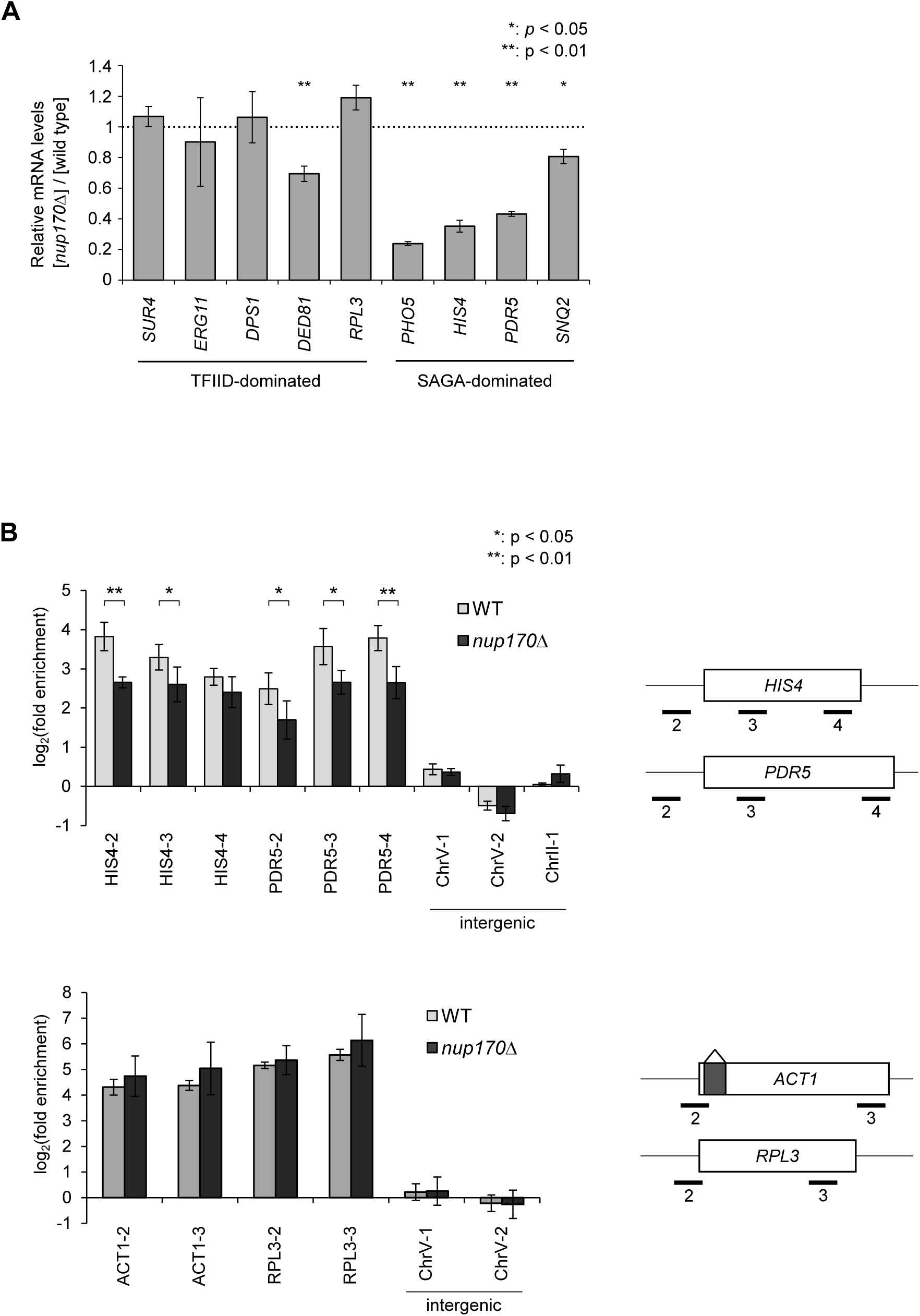
Loss of Nup170 preferentially suppresses expression of genes regulated by the SAGA coactivator complex. A) The fold change in mRNA levels of the indicated genes in the *nup170Δ* mutant compared with the wild-type strain were examined by RT-qPCR. The genes are categorized as either TFIID-or SAGA-dominated (Huisinga and Pugh, 2004; Barbaric et al., 2003). The bars indicate the standard deviation (*n* = 3). The genes whose mRNA level was significantly reduced in the *nup170Δ* mutant are marked with asterisks (Student’s *t*-test). B) The RNA Pol II density on various genes was examined by ChIP analysis using an anti-Rpb3 antibody. The enrichment of genomic regions was quantified by qPCR, and normalized with that of the intergenic regions (ChrV-2, ChrV-3, and ChrII-1). The regions amplified in the qPCR relative to the ORFs (indicated by boxes) are shown in panels to the right of the plots. The region 2 for each gene includes the transcription start site, and the regions 3 and 4 are located within the ORF. *ACT1* contains an intron indicated by a dark box. The regions showing significant difference in enrichment between the WT and the *nup170Δ* mutant, as determined by paired *t*-test, are marked with asterisks.

### Preinitiation complex association with SAGA-dominated gene promoters is defective in the *nup170Δ* mutant

To further understand the functional basis for the attenuated transcription of genes in the absence of Nup170, we examined the consequences of Nup170 loss on assembly of the preinitiation complex (PIC), including components of the SAGA complex, RNA Pol II, transcriptional activators, and histones, at promoter regions of several SAGA-dominated genes that show varying degrees of reduced expression in *nup170Δ* mutant: *SNQ2* (0.80 fold), *PDR5* (0.39 fold), and *HIS4* (0.35 fold) (Fig. 1A and Van de Vosse et al., 2013). ChIP analysis was employed to examine the association of Hfi1 (a SAGA complex protein), Rpb3, histone H3, and the transcriptional activators Pdr1 and Bas2 to the promoter regions of *PDR5, SNQ2*, and *HIS4* in the WT and *nup170Δ* mutant strains. As shown in Fig. 2B, in WT cells we detected the enrichment of Hfi1 and Rpb3 at the promoter regions of *PDR5, SNQ2*, and *HIS4*. Importantly, these interactions were significantly reduced in strains lacking Nup170 (Fig. 1B and 2B). We also examined the association of two transcriptional activators with the three target genes: Bas2 binding to *HIS4*, and Pdr1 binding to *PDR5* and *SNQ2*. We observed that, in the absence of Nup170, binding of Bas2 to the promoter region of *HIS4* was significantly reduced. Similarly, a smaller, but significant, reduction in the association of Pdr1 was detected with *PDR5* promoter region, while the binding of Pdr1 to *SNQ2* appeared unchanged (Fig. 2B).

**Figure 2.**
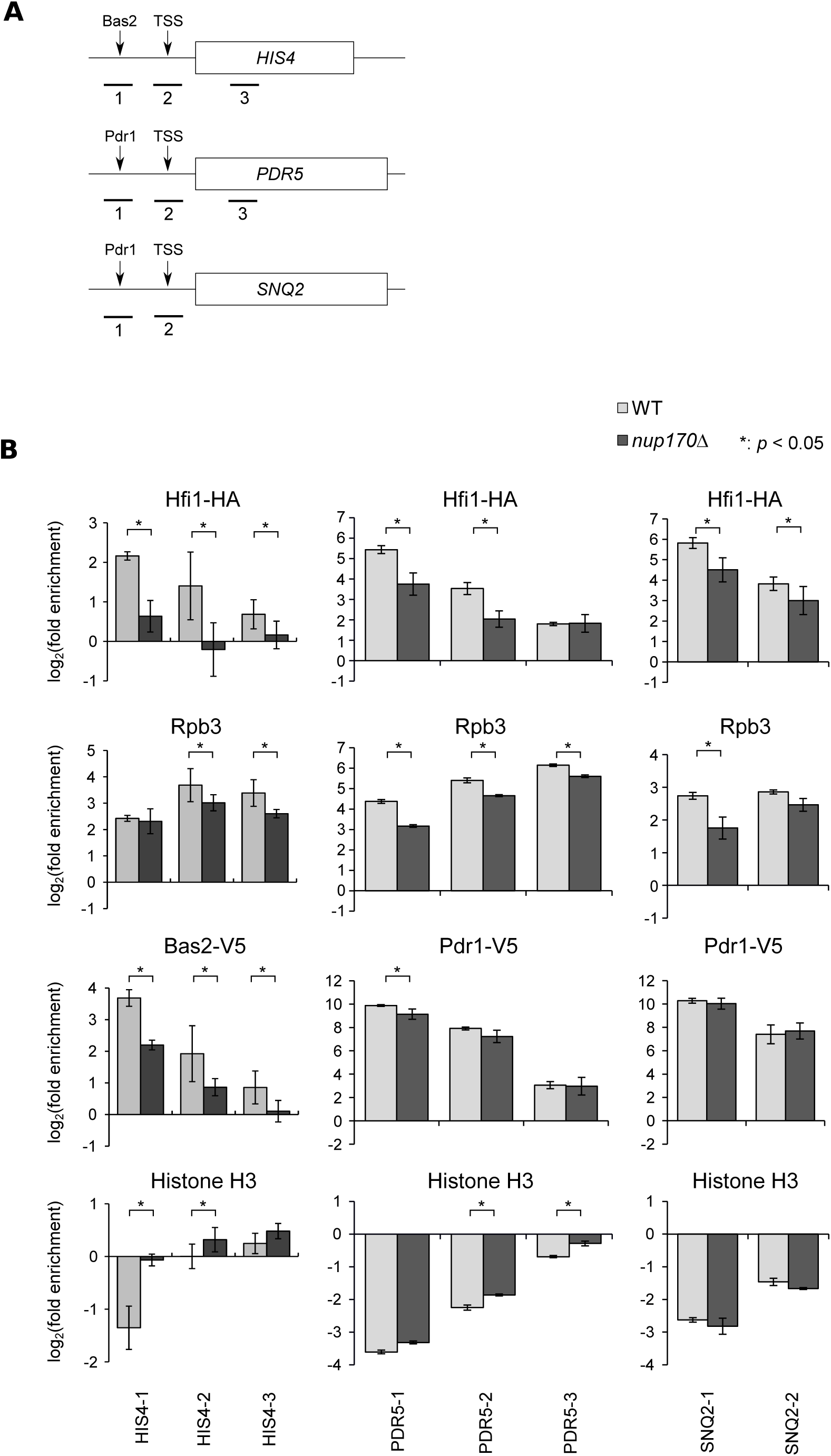
Genes repressed in the *nup170Δ* mutant exhibit reduced association of preinitiation complex components. A) ChIP analysis was performed using primer sets that detect the indicated regions of *HIS4, PDR5*, and *SNQ2*. The locations of these regions are shown in the schematic diagram. The region 1 for each gene includes the binding sites for the gene specific transcriptional activator (Bas2 or Pdr1). The region 2 includes the transcriptional start sites (TSS). The region 3 is located within the ORFs. B) WT and the *nup170Δ* mutant strains producing Hfi1-HA and either of Pdr1-V5 or Bas2-V5 were subjected to ChIP analysis using antibodies directed against the V5 or HA tags, Rpb3, and histone H3. The enrichment of the genomic regions was quantified using qPCR and normalized as described in Fig. 1B. The bars indicate standard deviation (*n* = 3). Regions showing significant differences in enrichment between the WT and the *nup170Δ* strain as determined using a paired *t*-test are marked with asterisks.

Histone association with the promoter regions also provide an indication of the transcriptional state of genes, as they are generally excluded from the promoter regions of transcribing genes (Hartley and Madhani, 2009). We confirmed this using ChIP analysis in the WT strain by showing that the promoter regions of *PDR5, SNQ2*, and *HIS4* were less enriched for histone H3 than the non-transcribed (background) regions (Fig. 2B). By contrast, in the *nup170Δ* mutant histone occupancy at the *HIS4* promoter region was significantly increased, consistent with its reduced transcription. For *PDR5* similar alterations were detected but were less robust, while *SNQ2* appeared unchanged (Fig. 2B). Notably, in the *nup170*Δ mutant the greater increases in histone occupancy associated with the *HIS4* and *PDR5* loci as compared to the *SNQ2* locus correlated with greater decreases in the expression of the *HIS4* and *PDR5* as compared to *SNQ2* (see Fig. 1A and Fig. 2B).

### Nup170 regulates cellular levels of the SAGA complex component Spt7

Our observations that numerous SAGA-dominated genes show reduced expression in the *nup170Δ* mutant lead us to investigate how the loss of Nup170 altered the SAGA complex. In order to examine the composition of the SAGA complex, we affinity purified the SAGA complex from WT and the *nup170Δ* mutant cells using a TAP-tagged version of the SAGA core component Spt20 (Spt20-TAP). Similar groups of proteins were isolated from WT and *nup170Δ* mutant cell extracts, and both displayed a banding pattern by PAGE similar to that previously reported for the Spt20-TAP-associated SAGA complex (see Spedale et al., 2010). However, a reproducible difference between WT and *nup170Δ* preparations was detected in the relative abundance of three protein species with apparent masses of ∼80 kD, ∼180 kD, and ∼200 kD (Fig. 3A). Specifically, Spt20-TAP associated proteins derived from the *nup170Δ* mutant showed reduced amounts of the 80 kD and 200 kD polypeptides, and increased levels of the 180 kD species (Fig. 3A). Previous analysis of the Spt20-TAP purified SAGA complex had identified the corresponding proteins as Spt8 (80 kD), Spt7 (200 kD), and a C-terminal truncation of Spt7 (180 kD) (Spedale et al., 2010). Consistent with these results, our mass spectrometry analysis of Spt20-TAP purified SAGA components identified Spt7 as the prominent species at 180 kD and 200 kD, and the presence of Spt8 at 80 kD (see Table S1). In regions of the gel containing the 80 kD and 200 kD proteins, lower levels of peptides derived from Tra1 were also detected, likely arising from degradation products of this ∼430 kD component of the SAGA complex (Table S1). The 200 kD species likely contains full-length Spt7, as peptides from the entire length of Spt7 were detected in this region of the gel. By contrast, the 180 kD species is likely a C-terminal truncation of Spt7, as no peptides after the amino-acid residue 938 where detected in this band (Table S2).

**Figure 3.**
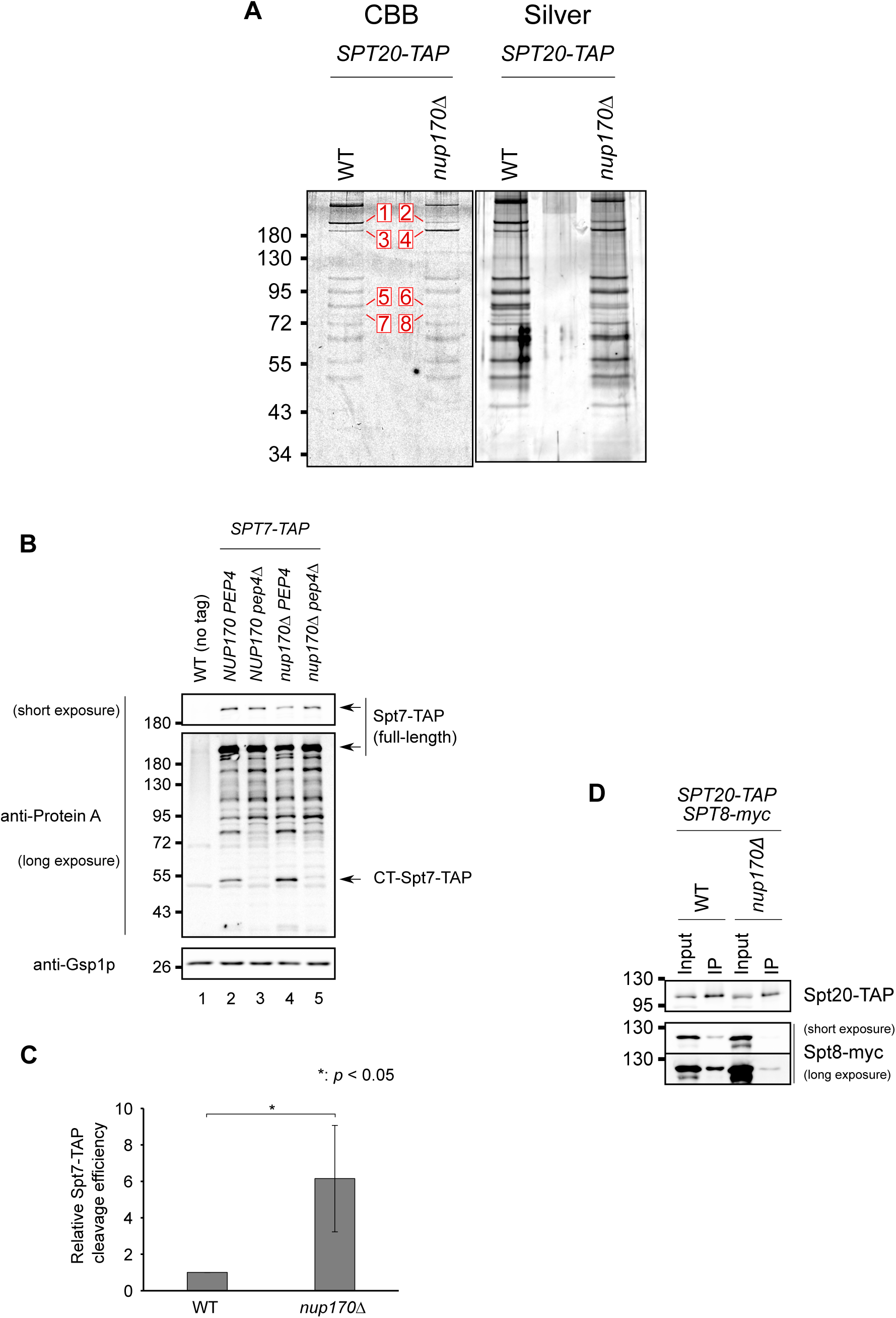
Nup170 is required for maintaining the cellular ratio of the SAGA and SLIK complexes. A) Spt20-TAP and associated proteins were purified from WT and *nup170Δ* cell lysates using IgG-conjugated Dynabeads. Protein complexes released from the beads were analyzed by the SDS-PAGE followed by the Coomassie brilliant blue (CBB) or the Silver staining. Protein species showing apparent differences in staining intensity between the WT and the *nup170Δ* mutant samples are indicated and numbered. The numbered protein species were cleaved and their peptides were analyzed by mass spectrometry. Results are shown in Table S1 and S2. B) The whole cell extracts were prepared from the indicated mutant cells producing Spt7-TAP, and extracts were analyzed by western blot using anti-Protein A and anti-Gsp1 (loading control) antibodies. The protein extracts from WT cells (no tag) were also analysed to reveal background signals. Indicated are full-length Spt7-TAP (two exposures are shown) and CT-Spt7-TAP (C-terminal fragment). C) The cleavage efficiency of Spt7-TAP in the wild-type (lane 2 in panel B) and the *nup170Δ* mutant (lane 4 in the panel B) were quantified. The ratio between the signal intensities of CT-Spt7-TAP (cleaved C-terminal fragment of Spt7) to full-length Spt7-TAP were measured in both strains. A relative Spt7-TAP cleavage efficiency was determined by dividing these ratios by that of the wild-type cells and resulting ratios were plotted. Thus, the ratio for the WT is one. The bars indicate the standard deviation (*n* = 4). The ratio values showed significant difference between the WT and the *nup170Δ* strain as determined using a paired *t*-test. D) Spt20-TAP and associated proteins were purified from WT and *nup170Δ* cells producing Spt8-myc. Total cell lysates (Input) and the purified protein complexes (IP) were analyzed by western blot using anti-Protein A and anti-myc antibodies. Two exposures are shown of the anti-myc antibody immunoblots. In all panels, the positions of mass markers are shown in kD.

The C-terminal truncation of Spt7 was previously shown to arise from the specific cleavage of Spt7 by vacuolar protease Pep4 following residue 1141 (Spedale et al., 2010; Mischerikow et al., 2009). To assess the role of Pep4 in the increased cleavage of Spt7 observed in the *nup170Δ* mutant, strains were constructed containing an Spt7-TAP fusion protein. In WT cells, full-length and proteolytic fragments of Spt7-TAP were visible by western blotting, including a polypeptide with molecule mass approximating the predicted 50 kD size of the Pep4-produced C-terminal Spt7-TAP fragment (CT-Spt7-TAP; Fig. 3B). Importantly, we observed that the amount of the CT-Spt7-TAP fragment was increased in the *nup170Δ* mutant (see Fig. 3B, lanes 2 and 4; Fig. 3C), and its presence in WT and the *nup170Δ* mutant was dependent on Pep4 (Fig. 3B; see *pep4*Δ and *nup170*Δ *pep4*Δ). Consistent with these results, we also observed a decrease in the amount of full length Spt7-TAP in the *nup170Δ* mutant, which was also Pep4-dependent. On the basis of these results, we concluded that the increased cleavage of Spt7 in the *nup170*Δ mutant is Pep4-dependent.

The C-terminal truncation of Spt7 is associated with a SAGA-like (SLIK) coactivator complex that lacks Spt8 (Pray-Grant et al., 2002; Sterner et al., 2002). Since core subunits such as Spt20 are components of both SAGA and SLIK, purification of Spt20-TAP reveals a ratio of full length Spt7 (∼200 kD) to its C-terminal truncation (∼180 kD) that reflects the in vivo ratio of the SAGA and SLIK complexes. In the *nup170Δ* mutant, the increased amount of the Spt7 C-terminal fragment bound to purified Spt20-TAP suggests both increased Pep4-dependent cleavage of Spt7 and a corresponding elevation in levels of SLIK. Consistent with this conclusion, our results revealed a reduction of Spt8-myc bound to Spt20-TAP purified from the *nup170Δ* mutant (Fig. 3D). These results suggest that Nup170 is required for maintaining the cellular ratio of the SAGA and SLIK complexes. This function for Nup170, however, does not appear to involve a direct interaction between Nup170 and SAGA or SLIK complex, as we did not detect Nup170 bound to the SAGA complex (data not shown).

### The NPC permeability barrier controls Pep4 accessibility to the nucleoplasm

Pep4 is a vacuolar protease that is processed to an active form in the lumen of the vacuole. Pep4 cleavage of Spt7 is presumed to occur following the movement of some portion of active Pep4 from the vacuole lumen to the nucleoplasm. While the mechanism by which Pep4 crosses the vacuolar membrane is undefined, it has been proposed that, once in the cytoplasm, Pep4 enters the nucleoplasm (Spedale et al., 2010). Pep4 does not contain a canonical nuclear localization signal, but its molecular mass (43 kD) would permit it to passively diffuse through NPCs and into the nucleoplasm (Timney et al., 2016). Importantly, NPCs lacking Nup170 exhibit significantly elevated permeability, with various reporter proteins (up to 126 kD) showing increased equilibration rates between the nucleoplasm and cytoplasm (Shulga et al., 2000; Timney et al., 2016). On the basis of these observations, we tested whether the loss of Nup170 would increase the accessibility of Pep4 to nuclear substrates. For these experiments, we constructed reporter substrates that are restricted to the cytoplasm or the nucleoplasm (Fig. 4A). The reporter proteins contain a segment of Spt7 (aa residues 1088-1180) that includes the Pep4 cleavage site (following residue 1141) flanked by GFP and GST. To this chimeric protein was added a nuclear localization (SV40 *NLS*) or nuclear export (PKI *NES*) signal, which drives the accumulate of the reporter protein in the nucleoplasm (GFP-*NLS*-tSpt7-GST) or cytoplasm (GFP-*NES*-tSpt7-GST) (Fig. 4B). Western blot analysis of WT cells producing these reporters revealed both the full-length (∼70 kD) and an ∼34 kD cleavage fragment predicted to be produced by Pep4 (Fig. 4C). Consistent with this conclusion, the presences of the ∼34 kD reporter cleavage fragments were dependent on Pep4 (Fig. 4C). Also of note, detection of the full-length reporters suggests the Pep4 substrates were not a limiting factor in the production of the ∼34 kD reporter cleavage fragments. We compared the levels of the NLS and NES reporter cleavage products (each normalized to their full-length counterpart) to produce a relative ratio of NLS to NES reporter cleavage in each strain. In WT cells this ratio was ∼ 0.6, suggesting cleavage of the nucleoplasmic NLS-containing reporter was less efficient than cleavage of the cytoplasm NES-containing reporter (see Fig. 4C, lanes 1 and 2; Fig. 4D). By contrast, in the *nup170*Δ strain, we observed a significant increase in the ratio of NLS to NES reporter cleavage products to ∼ 1.6 (Fig. 4D). These results are consistent with the conclusion that nuclear Pep4-dependent cleavage events are increased in the *nup170*Δ strain.

**Figure 4.**
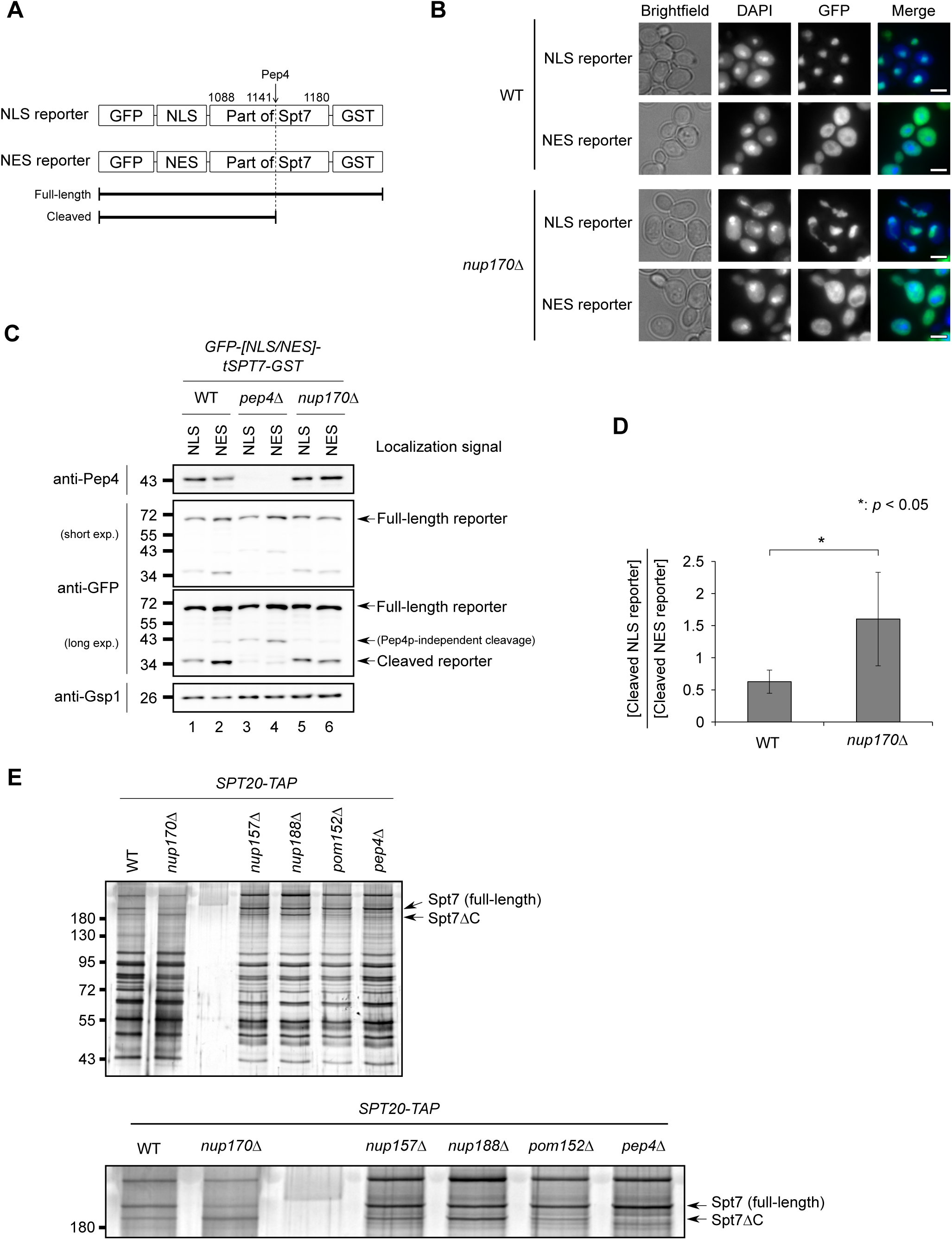
The loss of Nup170 allows increased accessibility of Pep4 to the nucleoplasm. A) Shown is a schematic diagram of the NLS and NES reporter proteins used for examining Pep4 activity in the cytoplasm and the nucleoplasm. The position of Pep4 cleavage site (following residue 1141) is indicated by an arrow. B) WT and the *nup170Δ* strains containing the reporter gene constructs were grown in YP raffinose media supplemented with 2% galactose for 3 h to induce production of the NLS- and NES-reporter proteins depicted in panel A. The cells were fixed with 5% formaldehyde and examined using an epifluorescence microscope. Images showing the positions of cells (Brightfield), the nuclei (DAPI), reporter proteins (GFP), and merged images of DAPI and GFP (Merge) are shown. Bars, 5 μm. C) WT, *pep4Δ*, and *nup170Δ* strains containing the reporter gene constructs were induced to produce the nuclear GFP-*NLS*-tSpt7-GST (NLS) or cytoplasm GFP-*NES*-tSpt7-GST (NES) reporter protein. Cell lysates were analyzed by western blotting using the indicated antibodies to detect Pep4, the full length and cleaved NLS (lanes 1,3,5) and NES (lanes 2,4,6) reporter proteins (anti-GFP), and a loading control (Gsp1). Two exposures are shown of the anti-GFP immunoblots. In all panels, the positions of mass markers are shown in kD. D) The difference in efficiency of Pep4-dependent cleavage of the GFP-*NLS*-tSpt7-GST and GFP-*NES*-tSpt7-GST reporter proteins between WT and the *nup170Δ* strain were compared. The cleavage efficiencies of the reporter proteins were measured as the ratio between the signal intensities of cleaved reporters and the corresponding full-length proteins. A relative cleavage efficiency of the NLS reporter to that of the NES reporter was then calculated, and the resulting ratio was plotted. The bars indicate the standard deviation (*n* = 3). The ratio values show significant differences between WT and *nup170Δ* strains as determined using a Student’s *t*-test. E) Spt20-TAP was purified from lysates of WT and the indicated mutant strains. Purified Spt20-TAP-containing complexes were analyzed by the SDS-PAGE followed by silver staining. Two bands, the full-length Spt7 and the C-terminally truncated Spt7 (Spt7*Δ*C), are indicated by arrows. The image at the bottom to shows a magnification of the Spt7 and Spt7*Δ*C containing-region of the top image. In all panels, the positions of mass markers are shown in kD.

We propose that the increased cleavage of endogenous Spt7 and the NLS-tSpt7 reporter in the *nup170*Δ stain occurs as a consequence of increased diffusion of Pep4 into the nucleoplasm. To further test this idea, we examined whether loss of another Nup, Nup188, also previously shown to contribute to the NPC diffusion channel (Shulga et al., 2000), would similarly increase the cleavage of endogenous Spt7. For these experiments, Spt20-TAP was introduced into a *nup188Δ* mutant as well as the two mutant strains lacking either Nup157 (a paralog of Nup170) or Pom152, two Nups not required for maintaining the passive diffusion barrier of the NPC (Shulga et al., 2000). Spt20-TAP was then purified from these mutant strains and the levels of associated full-length Spt7 and its Pep4-dependent C-terminal truncation were compared. In the *nup157*Δ, and *pom152*Δ mutants, the levels of Spt7 and its C-terminal truncation were similar to that detected in WT cells (Fig. 4E). However, like the *nup170Δ* mutant, the amount of the truncated Spt7 was increased in the *nup188Δ* mutant (Fig. 4E). These results support the conclusion that passive diffusion channels of the NPCs control accessibility of Pep4 to Spt7 and thus levels of the SLIK complex.

The passive diffusion properties of yeast NPCs are also altered by certain stress conditions induced by treatment of the cells with hydrogen peroxide or 1, 6-hexanediol (Mason et al., 2005; Shulga and Goldfarb, 2003). Therefore, we tested the impact of treating cells with these compounds on the levels of SAGA and SLIK complexes, again comparing levels of full length and truncation Spt7 as a measure of the relative levels of these complexes. As shown in Fig. 5, treatment of cells with various concentrations of hydrogen peroxide or 1, 6-hexanediol led to the increased cellular levels of the Spt7 truncation relative to full length Spt7. Thus, similar to strains lacking Nup170 or Nup188, stress conditions previously shown to increase the size limits of the NPC passive diffusion channel also induce greater cleavage of Spt7 and an increase in the SLIK:SAGA complex ratio.

**Figure 5.**
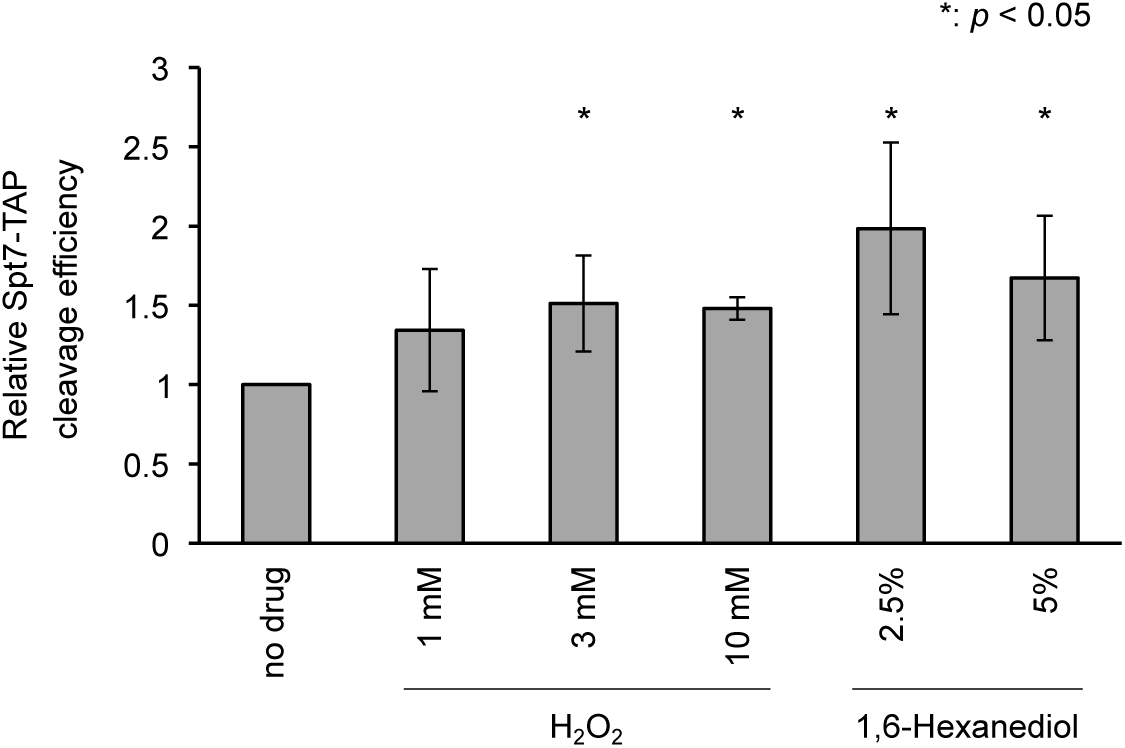
Cell stress can induce increased Spt7 cleavage. Cells expressing *SPT7-TAP* were grown in YPD medium and 1 h prior to harvesting cells were treated with or without hydrogen peroxide or 1, 6-hexanediol at the indicated concentrations. Cell lysates were analyzed by western blotting using anti-Protein A (to detect Spt7-TAP and the cleaved CT-Spt7-TAP) and anti-Gsp1 antibodies (loading control). The ratio between the signal intensities of CT-Spt7-TAP (cleaved C-terminal fragment of Spt7) to full-length Spt7-TAP were measured for each of the various conditions. Spt7-TAP cleavage efficiencies under the various conditions relative to the no drug control where determined as described in Fig. 3C. The bars indicate the standard deviation (*n* = 3). The treatment conditions where the Spt7 cleavage was significantly increased compared to the “no drug” control, as determined by a paired *t*-test, are indicated by asterisks.

**Figure 6.**
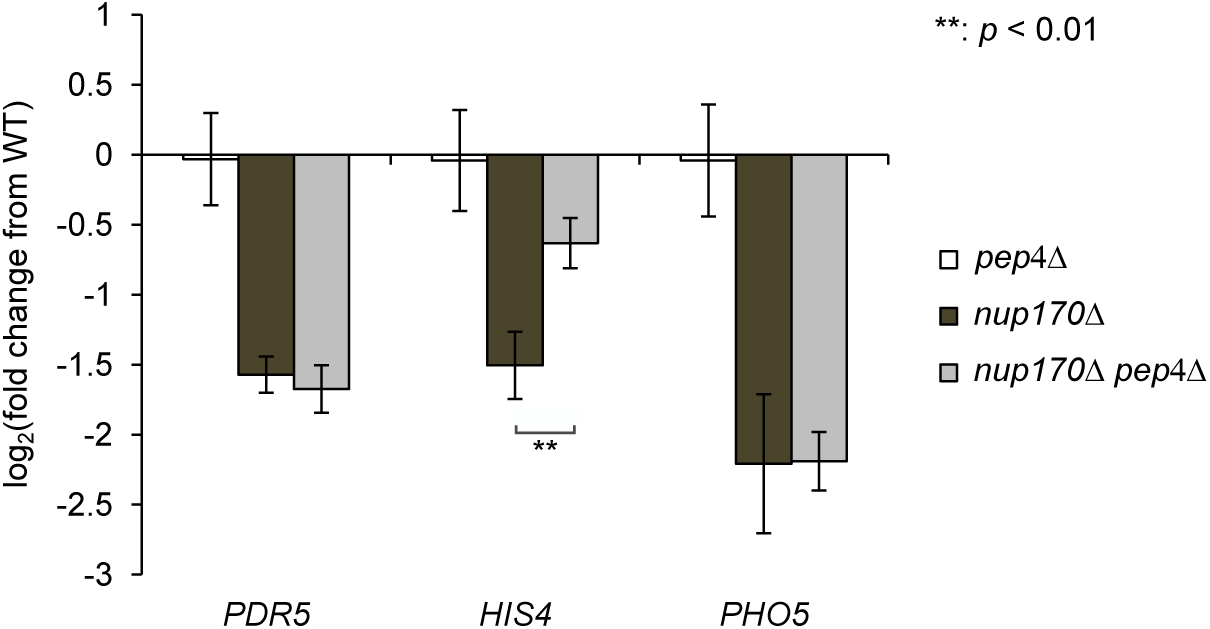
Reduced expression of *HIS4* observed in the *nup170*Δ mutant is dependent on Pep4. Total RNA was isolated from WT, *nup170Δ, pep4Δ* and *nup170Δ pep4Δ* mutant strains, and the mRNA levels of the indicated genes were examined by the RT-qPCR. Levels of mRNA encoded by the indicated genes in each mutant were normalized to amounts in the WT strain. Fold changes in amounts detected in the mutants relative to WT cells were plotted. The bars indicate the standard deviation (*n* = 3). Significant differences in mRNA levels between the *nup170Δ* and the *nup170Δ pep4Δ* strains, as determined by a Student’s *t*-test, are indicated by asterisks.

### Preventing Spt7 cleavage in the *nup170Δ* background partially reversed the reduced expression of the SAGA-dominated genes

On the basis of our analysis of the *nup170Δ* mutant, we predict that changes in cellular levels of the SAGA and SLIK complexes contribute to some of the observed alterations in transcription detected in this mutant. In order to investigate the effect of altering SAGA-SLIK balance on the gene expression pattern, we prevented the SLIK complex formation in the *nup170Δ* mutant by disrupting the *PEP4* gene, and we examined the effect of the *pep4*Δ mutation on the expression of three SAGA-dominated genes, *PHO5, PDR5*, and *HIS4* (see Fig. 2). As shown in Fig. 5, the *pep4Δ* mutation suppressed the reduction of *HIS4* mRNA levels seen in the *nup170Δ* background, while the mRNA level of *PDR5* and *PHO5* was not altered by *PEP4* deletion. These results support the conclusion that at least a subset of the transcriptional reductions observed in the *nup170Δ* mutant are linked to Pep4 cleavage of Spt7.

## Discussion

Transcriptome analysis of *nup170*Δ mutants revealed that Nup170 is required to repress the transcription of certain genes (Van De Vosse et al., 2013) while supporting the transcription of other genes (Van De Vosse et al., 2013; Fig. S1). These observations implicate Nup170 in distinct regulatory processes. In examining those genes that show reduced levels of transcripts in *nup170*Δ mutants, we observed that many are regulated by the SAGA transcriptional activator. Consistent with this, analyses of a subset of these genes in the *nup170*Δ mutant revealed reduced levels of bound preinitiation complex (PIC), including components of the SAGA complex, transcriptional activators, and RNA Pol II. Concomitant with these changes are alterations in the relative cellular levels of the SAGA and SLIK complexes, with increased levels of the SLIK complex detected in the *nup170*Δ mutants. Nup170 does not appear to directly interact with SAGA/SLIK complexes, but rather its role in mediating levels of these complexes is indirect and linked to changes in passive diffusion barrier of the NPC. These results lead us to propose that changes in passive diffusion rates through NPCs, as would occur during periods of altered cell physiology (e.g. stress) represents a mechanism for globally modulating nuclear functions including transcriptional processes such as those regulated by the SAGA/SLIK complexes.

Nups can influence gene expression by various mechanisms, including through their direct interactions with gene loci and the transcriptional machinery, or indirectly through their roles in regulating nuclear transport of transcriptional regulators. Some Nups are envisaged to contribute to multiple process, such as mammalian Nup98, which binds to and regulates the activity of nuclear transcription coactivators (Kasper et al., 1999) and RNA helicases (Capitanio et al., 2017, 2018) while also functioning in nuclear transport at NPCs (Powers et al., 1997; Oka et al., 2010). Transcriptional changes arising from mutations in such a multi-functional Nup are, therefore, likely to be a product of functional alterations in the various pathways that they contribute to, and their cumulatively impact on gene expression. Our analysis of the *nup170*Δ mutant suggests that this Nup can also influence transcription by multiple mechanisms. Previous studies showed that Nup170 associates with subtelomeric chromatin and is required for the assembly of chromatin-associated silencing complexes containing Sir4 (Van De Vosse et al., 2013). In this instance, the loss of Nup170 reduced silencing of numerous subtelomeric genes leading to their increased expression. Notably, recent studies showed that this function of Nup170 is likely performed by Nup170 present in a structure distinct from NPCs, termed the Snup-complex, containing a subset of Nups and silencing factor Sir4 (Lapetina et al., 2017).

In addition to its role in promoting gene silencing, here we report that Nup170 also supports active transcription of a subset of genes, most regulated by the SAGA complex (Fig. S1). Multiple observations lead us to conclude that Nup170 supports transcription indirectly. In support of this idea, we failed to detect physical interactions between Nup170 and SAGA complex components or members of the downregulated group of genes using IP or ChIP analysis (data not shown). Despite this, in cells lacking Nup170 the SAGA-regulated genes examined showed reduced levels of bound SAGA complex (Fig. 2B).

Consistent with the reduced levels of SAGA complex associated with genes normally regulated by this complex (Fig. 2), we also detected in the *nup170*Δ mutant reduced cellular levels of SAGA complex relative to the related SLIK complex (Fig. 3). A clue to how Nup170 might influence the ratio of SAGA to SLIK complexes came from an examination of the mechanism of SLIK complex formation. Previous studies have shown that the SLIK complex arises from the cleavage of the SAGA complex component Spt7 by the vacuolar protease Pep4 (Spedale et al., 2010). The resulting truncated Spt7 fails to bind Spt8, the lack of which is a defining feature of the SLIK complex (Pray-Grant et al., 2002; Sterner et al., 2002). Of note, Spt8 is one of two SAGA components (along with Spt3) that interacts directly with the TATA binding protein (TBP) (Han et al., 2014). As a consequence, the SLIK complex lacking Spt8 exhibits reduced binding to TBP (Sterner et al., 1999). This is consistent with our observation that the enrichment of Hfi1 (a common component of SAGA and SLIK) at the promoter regions of the SAGA-dependent genes examined was significantly reduced in the *nup170Δ* mutant (Fig. 2B).

We showed that the increased levels of the SLIK complex detected in the *nup170*Δ mutant were dependent on Pep4, leading us to conclude that Nup170 was functioning to upregulate Pep4-mediated cleavage of Spt7. These data are consistent with previous literature identifying Pep4 as the protease responsible to cleavage of Spt7 and production of the SLIK complex (Spedale et al., 2010). Curiously, Pep4 is generally considered a luminal vacuolar protein. It contains a signal sequence that directs prepro-Pep4 to and across the ER membrane and into the ER lumen. From here Pep4 is transported to the vacuole lumen. It is here that a vacuolar protease activates Pep4 by removing its N-terminal propeptide (van den Hazel et al., 1996). Thus, for Pep4 to cleave Spt7, which is localized to the nuclear interior (Spedale et al., 2010), it is presumed that some portion of Pep4 must exit vacuoles and enter the cytoplasmic compartment. How this may occur is unknown, however this is not an isolated phenomenon as other examples of vacuolar/lysosomal proteins functioning in the cytoplasmic/nuclear compartments have been reported, especially under stress conditions. For example, the endopolyphosphatase Ppn1 (Phm5) is processed to its enzymatically active, mature form in vacuoles but active Ppn1 is detected in the cytoplasm (Lichko et al., 2004; Shi and Kornberg, 2005; Reggiori and Pelham, 2001). In addition, various cytoplasmic tRNAs are cleaved by a vacuolar nuclease Rny1 (Thompson et al., 2008; Thompson and Parker, 2009). Nuclear activities of metazoan lysosomal enzymes have also been reported. The lysosomal protease cathepsin L is released from the lysosome and reaches the nucleus (Tholen et al., 2014) where it cleaves the CDP/Cux transcription factor to promote S phase progression (Goulet et al., 2004) and histone H3 during mouse embryonic stem cell differentiation (Duncan et al., 2008).

Our data also support the conclusion that Pep4 exits vacuoles and enters the cytoplasm and the nucleus. Using reporter substrates containing the Pep4 cleavage site of Spt7 and either an NES or NLS, we were able to position the Pep4 substrate in the cytoplasm or the nucleus and assay for their cleavage. Both substrates exhibited Pep4-dependent cleavage in WT and *nup170*Δ cells (Fig. 4). These results supported the conclusion that Pep4 can enter the nucleus where it would have access to endogenous Spt7. Importantly, in the *nup170*Δ mutant, cleavage of the nuclear NLS-containing reporter was increased, while cleavage of the cytoplasmic NES-containing reporter appeared reduced, leading to an increase in the ratio of the NLS-reporter:NES-reporter cleavage products (Fig. 4D). These data are consistent with the higher levels of the Spt7-truncation detected in the *nup170*Δ mutant (Fig. 3A, 3B and 3C), and they suggest this phenotype arises from increased levels Pep4 in the nucleus in the *nup170*Δ mutant. Of note, attaching a tag (GFP and HA) to Pep4 for the purpose of examining its localization was not useful as these tagged forms of Pep4 appeared to be unstable and their cellular levels were severely reduced relative to untagged Pep4. Consistent with this observation, cells producing Pep4-GFP showed reduced cleavage of Spt7(data not shown).

We were posed with the question of how Pep4, once released from vacuoles, would enter the nucleoplasm, and why entry is more efficient in *nup170*Δ cells. We concluded that Pep4 entry into the nucleus was unlikely to be mediated by NTFs. First, Pep4 lacks any detachable nuclear localization signals within its primary sequence that would facilitate nuclear import. Second, the increase in nuclear Pep4 activity detected in the *nup170*Δ mutant was also unlikely to be explained by increased NTF-mediated nuclear import as transport changes of this type have not been observed in the *nup170*Δ mutant (Shulga et al., 2000). Instead, an alternative explanation is that movement of Pep4 across the NE and into the nucleus is controlled by the passive diffusion barrier of NPCs. Consistent with this idea, previous studies have shown that sieving limit for the NPC diffusion channel is greater in cells lacking Nup170 (Shulga et al., 2000; Timney et al., 2016). Shulga et al., also showed an increase in the size of the NPC diffusion channel in cells lacking Nup188, a binding partner of Nup170 within the inner ring complex of the NPC. By contrast, the loss of two other Nup170 binding partners, Nup157 and Pom152, does not alter the diffuse rates through NPCs (Shulga et al., 2000). Consistent with the roles of these various Nups in passive diffusion, the levels of the Spt7 truncations (i.e. Pep4 cleavage) were increased in the *nup170*Δ and *nup188*Δ mutants, while mutants that did not affect diffusion (*nup157*Δ and *pom152*Δ) contained WT levels of the Spt7 truncation (Fig. 4E). Moreover, in comparing the *nup170*Δ and *nup188*Δ mutants, we observed that the higher sieving limits reported for NPCs lacking Nup170 versus those lacking Nup188 (Shulga et al., 2000) directly correlated with higher levels of truncated Spt7 in the *nup170*Δ mutant as compared to *nup188*Δ mutant (Fig. 4E).

Cumulatively, the results presented in this manuscript suggest that the properties of the passive diffusion channel of the NPC can function to mediate levels of transcriptional complexes in the nucleoplasm. Specifically, our study revealed that the relative abundance of the SAGA and SLIK complexes are influenced by Pep4 and the passive diffusion properties of the NPC. We envisage that there are likely other examples of factors controlling nuclear functions whose entry into the nucleus are mediated by the NPC passive diffusion barrier rather than facilitated NTF-mediated transport. Moreover, this may include a broader spectrum of cytoplasm molecules than previous considered. This is emphasized by recent work from Timney et al (2016) suggesting that the NPC does not impose strict passive diffusion limit but rather it functions as a soft barrier that gradually restricts the movement of molecules as they increase in mass. These observations suggest that molecules of greater mass than previously anticipated can passively diffuse across the NE, albeit with decrease rates as their size increases. Consistent with this idea, many proteins and protein complexes of up to ∼100 kD (and lacking nuclear transport signals) appear to partition equally between the cytoplasm and the nucleoplasm (Wühr et al., 2015).

Our observations that specific transcriptional events, such as those controlled by the SAGA and SLIK complexes, are linked to the state of the passive diffusion barrier suggest that conditions that alter its properties, such as stress, represent a potential control point for regulating transcription and other nuclear functions. In support of such a mechanism, previous studies have shown that stressing yeast cells using hydrogen peroxide induces an increase in the size of the NPC passive diffusion channel (Mason et al., 2005), and, as shown in Fig. 5, an increase in levels of the Spt7 truncation and the SLIK complex. Of note, cellular responses to other stress conditions have also been linked to the activity of the SLIK complex (Spedale et al., 2010). For example, mutant cells in which the SLIK complex predominates are more resistant to the growth inhibitory effects of rapamycin treatment than WT cells, while cells in which SAGA is predominate are more sensitive to rapamycin than WT (Spedale et al., 2010).

Our observations that transcriptional changes in the *nup170*Δ mutant arise from the loss of several distinct functions of this Nup suggests that one must generally consider the multifunctional potential of any Nup, including a role in the passive diffusion barrier, when assessing its roles in nuclear functions, such as transcriptional regulation. With this in mind, it will be of future interest to assess the similarities and differences in the transcriptome profiles of various nups mutants that alter the passive diffusion barrier. This may provide clues as to the broader effects of altering the passive diffusion barrier on transcription, and whether specific Nups, through their signature impacts on passive diffusion, may influence specific transcriptional events. We speculate that physiological or environmental changes that alter the function of Nups forming the passive diffusion barrier, either through their modification or degradation, could represent a mechanism for transiently regulating transcriptional, for example during periods of stress (as discussed above) or in response changes in growth, cell shape, or during developmental stages when the passive diffusion barrier also changes (Feldherr, 2004; Jiang and Schindler, 2004; García-González et al., 2018). In addition, continuous and irreversible changes in the permeability of NPCs have been proposed to occur in postmitotic metazoan cells as they age (D’Angelo et al., 2009). This has been suggested to arise as a consequence of accumulated damage to long-lived Nups, including Nup155, the counterpart of Nup170 (D’Angelo et al., 2009; Savas et al., 2012). The consequences of these age-related changes in the passive diffusion barrier of the NPC remain to be determined, but it seems reasonable to speculate that many nuclear functions are impacted, including transcription. Similarly, it is important to consider whether disease states linked to mutations in the human counterparts of several yeast Nups, such as Nup170, Nup188, and Nup116 (Zhang et al., 2008; Simon and Rout, 2014; Haskell et al., 2017; Beck and Hurt, 2017), occur as a consequence of constitutive defects in the passive diffusion barrier.

## Materials and Methods

### Yeast strains and cultures

Yeast strains used in this study are shown in Table S3. All strains were grown in YP (1 % yeast extract, 2% Peptone) plus appropriate carbon source (2% glucose or 2% raffinose) at 30°C. For the gene induction under control of *GAL1* promoter, galactose (from a 20% stock solution) is added directly into the culture media to a final concentration of 2%. For stress treatment with hydrogen peroxide and 1,6-hexanediol, the chemicals from stock solutions (100 mM hydrogen peroxide and 50% 1,6-hexanediol in water) were directly added to the media.

### Western blot

For analyzing the whole cell lysate of yeast in normal growing conditions, cells were grown in the YP glucose medium at 30°C to an OD_600_ of ∼ 1.0. For preparing yeast cells after the galactose induction, cells were grown in the YP raffinose medium to OD_600_ of ∼ 0.5, followed by incubation with an additional 2% galactose for 3 h. One OD_600_ cells were collected and resuspended in 100 μl of SDS-PAGE sample buffer, followed by brief sonication and heat denaturation at 65°C for 10 min. The protein samples were separated by SDS-PAGE and transferred on the nitrocellulose membranes. Proteins were detected by Western blotting using anti-Protein A from rabbit (for detecting TAP fusion proteins; P3775; Sigma), anti-Gsp1 from rabbit (Makhnevych et al., 2003), anti-Pep4 from rabbit (a gift from Dr. Gary Eitzen (University of Alberta)) and anti-GFP from mouse (clones 7.1 and 13.1; SKU 11814460001; Roche), and secondary antibodies, HRP-conjugated anti-rabbit IgG and HRP-conjugated anti-mouse IgG (BioRad). The protein signals were visualized using ECL Western Blotting Detection Reagent (GE), and scanned using an ImageQuant LAS 4000 (GE) imaging system.

### The mRNA extraction and quantitative RT PCR

The total RNA from yeast was extracted by the hot phenol method (Van De Vosse et al., 2013). Ten ml of yeast culture was grown to an OD_600_ of 0.8 – 1.0. Cells derived from 5 OD_600_ of cell culture were collected and resuspended in 0.5 mL TES (10 mM Tris-HCl pH 7.5, 1 mM EDTA, 100 mM NaCl). The suspension was well mixed with 0.5 mL water-saturated phenol using vortex. Then the sample was incubated for 1 h at 65°C in order to extract RNA from the cells. After centrifugation, the water layer containing RNA was further purified by mixing with phenol/chloroform/isoamyl alcohol (25:24:1). The RNA was precipitated with ethanol, dried, and dissolved in DEPC water. Two micro grams of total RNA was subjected to the reverse transcription using the Superscript II kit (Life Technologies). The relative gene abundance in the cDNA sample was examined by the quantitative PCR (qPCR) experiment on MX3000 (Agilent) using PerfeCTa SYBR green PCR mix (Quanta Bioscience). Oligonucleotides for the reaction were summarized in Table S4. The change in the mRNA level of the genes among various mutants were evaluated by the ΔΔC_T_ method (Livak and Schmittgen, 2001). The C_T_ value for each gene target was normalized against the internal control (*ACT1* and *TUB1*) to give the ΔC_T_ value. The ΔΔC_T_ was calculated as a difference in ΔC_T_ values between the mutant and the wild type strain. The change in the mRNA level of the genes among the mutant was given as 2^−ΔΔCT^, based on the assumption that amplification efficiency of the PCR reaction is 100%.

### Chromatin immunoprecipitation (ChIP)

ChIP experiment was performed as described previously (Hecht and Grunstein, 1999; Van De Vosse et al., 2013). Fifty ml of culture at an OD_600_ of 0.8-1.0 was subjected to crosslinking with 4% formaldehyde for 20 min at room temperature. The crosslinking reaction was quenched by incubation with 125 mM glycine for 5 min. Cells were resuspended in lysis buffer (50 mM HEPES-KOH, pH 7.5, 140 mM NaCl, 1 mM EDTA, 1% Triton X-100, 0.1% sodium deoxycholate) and disrupted by vortex in the presence of glass beads, and the chromatin DNA was sheared by sonication (Branson Sonifer 250) to an average fragment size of 300-500 bp. The lysate was incubated with 3 μg of antibodies (either of anti-V5 (SV5-Pk1; ab27671; Abcam), anti-HA (12C5; 11583816001; Roche), anti-Rpb3p (1Y26 (1Y27); ab81859; Abcam), anti-Histone H3 (ab46765; Abcam)) and 50 μl suspension (1.5 mg of beads) of Protein G-conjugated Dynabeads (Life Technologies) for 2 h at 4°C. Chromatin regions captured by Dynabeads were collected with magnetic stands. After washing, crosslinks were reversed by incubation at 65°C overnight.

The enrichment of chromatin regions was examined by qPCR on MX3000 (Agilent) using PerfeCTa SYBR green PCR mix (Quanta Biosciences). Oligonucleotides used for the qPCR were summarized in Table S4. Relative enrichment of the gene loci immunoprecipitated with the protein of interest was evaluated by the ΔΔC_T_ method (Livak and Schmittgen, 2001). The C_T_ values for the amplicon from the IP sample were firstly normalized against the corresponding input to give the ΔC_T_ values. The ΔΔC_T_ value was calculated as a difference in ΔC_T_ values between the gene locus of interest and the non-transcribed control (ChrV-1, ChrV-2 and ChrII-1). These non-transcribed controls are located in the middle of a > 1 kbp region without any observed transcription (ORFs, SUTs, CUTs, or other RNA polymerase-dependent transcription) (Xu et al., 2009). The relative enrichment of the specific genomic loci over background was given as 2^−ΔΔCT^, based on the assumption that amplification efficiency of the PCR reaction is 100%.

### Immunoisolation of the SAGA (and SLIK) complex and mass spectrometry analysis

Cells derived from a one liter culture (OD_600_ of 1.0) of a strain expressing *SPT20-TAP* were collected and flash frozen with a liquid nitrogen. Cell samples were subjected to cryogrinding (Retsch PM100) to produce frozen lysate powder (Oeffinger et al., 2007). One gram of the lysate powder was resuspended in the IP buffer (20 mM HEPES-KOH pH 7.5, 110 mM potassium acetate, 100 mM NaCl, 2 mM MgCl_2_, 0.1 % Tween 20, 5% glycerol) plus cOmplete protease inhibitor cocktail without EDTA (Roche). The suspension was clarified by a series of consecutive centrifugations (500 × *g* for 10 min, 20000 × *g* for 20 min, and 100000 × *g* for 1 h) and the supernatant was collected. The cleared lysate was incubated with 3 mg IgG-conjugated Dynabeads for 1 h at 4°C. The beads were collected using a magnetic stand and washed 5 times with the IP buffer (without protease inhibitor cocktail). The protein complexes bound to the beads were released by a treatment with TEV protease (kindly provided from Dr. Ben Montpetit, University of California Davis) for 1 h at RT. Proteins released from the beads were precipitated with 10 % TCA, and resuspended in SDS-PAGE sample buffer.

Isolated proteins were separated on the SDS-PAGE gels and visualized by either CBB, silver staining, or western blotting using anti-Protein A from rabbit (for detecting TAP fusion proteins; P3775; Sigma), anti-myc from mouse (9E10; 11667149001; Roche) and anti-HA from mouse (12CA5; 11583816001; Roche), and secondary antibodies, HRP-conjugated anti-rabbit IgG and HRP-conjugated anti-mouse IgG (BioRad). For mass spectrometry analysis, the protein samples in the CBB-stained gels were excised and submitted to the Alberta Proteomics and Mass Spectrometry Facility (Department of Biochemistry, University of Alberta) for identification. The data were analyzed with the software Proteome Discoverer (Thermo).

### Microscopy

Cells producing the indicated GFP fusion proteins were grown to an OD_600_ of ∼ 1.0, and fixed with 5% formaldehyde in 100 mM potassium phosphate, pH 6.5 for 10 min. The fixed cells were washed with PBS twice and resuspended in a mounting medium (Dapi Fluoromount-G; Southern Biotech). The samples were put on microscope slides, and the images were observed on an AxioObserver.Z1 microscope (Zeiss) equipped with an UPlanS-Apochromat 100×/1.40-NA oil objective lens (Zeiss) and an AxioCam MRm digital camera with a charge-coupled device (Zeiss). Image manipulations (crop and normalization) were performed on the software ImageJ (NIH).

## Acknowledgements

We thank Dr. Ben Monpetit (University of California, Davis) and Dr. Gary Eitzen (University of Alberta) for providing the indicated reagents. We also thank B. Montpetit for critical reading of the manuscript. Funding for this work is supported by Canadian Institutes of Health Research (MOP 106502 and 36519). The authors declare no competing financial interests.

