## Supplemental Tables and Figures for "Passive diffusion through nuclear pore complexes regulates levels of the yeast SAGA and SLIK coactivators complexes"

**Table S1. Yeast proteins identified by the mass spectrometry analysis of the protein species highlighted in Figure 3A.**

| Sample No. | Gene Name | Score | Coverage | # PSMs | MW [kD] |
|---|---|---|---|---|---|
| 1 | SPT7 | 35.5 | 12.3 | 13 | 152.5 |
| 2 | TRA1 | 8.9 | 1.3 | 4 | 432.9 |
| | SPT7 | 7.6 | 2.9 | 3 | 152.5 |
| 3 | SPT7 | 15.9 | 6.2 | 6 | 152.5 |
| | TRA1 | 4.6 | 0.7 | 2 | 432.9 |
| 4 | SPT7 | 51.4 | 16.1 | 23 | 152.5 |
| | TRA1 | 13.3 | 1.8 | 6 | 432.9 |
| 5 | SGF73 | 26.9 | 18.3 | 16 | 72.8 |
| | SPT8 | 20.7 | 14.6 | 7 | 66.1 |
| | SPT20 | 5.7 | 5.0 | 2 | 67.8 |
| 6 | SGF73 | 14.5 | 6.9 | 7 | 72.8 |
| 7 | SSA2 | 22.0 | 16.1 | 9 | 69.4 |
| | SSB1 | 8.4 | 7.8 | 4 | 66.6 |
| 8 | SGF73 | 7.8 | 4.6 | 3 | 72.8 |

Protein samples derived from the numbered species in Fig. 3A were submitted to the mass spectrometry analysis and the identified gene products in each sample are summarized. From the analysis with the software Proteome Discoverer, the score, the percentage of protein sequence coverage by the identified peptides (Coverage), the total number of peptide spectral matches (PSM), and the calculated molecular weight (MW) for each gene product were determined and are shown here.

**Table S2. Peptides derived from Spt7 identified in the mass spectrometry analysis.**

| Peptide Sequences | position | Sample 1 | Sample 2 | Sample 3 | Sample 4 |
|---|---|---|---|---|---|
| GNIALNVEK | 65-73 | N/D | N/D | N/D | 1 |
| SDDVSSQTIK | 95-104 | N/D | N/D | N/D | 2 |
| FAEDEDYDDEDENYDEDSTDVK | 232-253 | 2 | N/D | 1 | 2 |
| NLDSISSSNIEIDDER | 261-276 | 1 | N/D | N/D | 1 |
| TNNVEEImGNWNK | 294-306 | 1 | N/D | N/D | 1 |
| SDLEAATDEQDRENTNDEPDTNQK | 341-364 | N/D | N/D | 1 | 1 |
| HLLSSIQQK | 401-409 | N/D | N/D | N/D | 1 |
| KSQLGISDYELK | 410-421 | N/D | N/D | N/D | 1 |
| SQLGISDYELK | 411-421 | 1 | N/D | N/D | 1 |
| IGQEELYEAcEK | 440-451 | 1 | 1 | 1 | 1 |
| NYTEHSTPFLNK | 458-469 | 1 | N/D | N/D | 1 |
| SmDLNTVLK | 485-493 | N/D | N/D | N/D | 1 |
| SMDLNTVLK | 485-493 | N/D | N/D | N/D | 1 |
| TASSTVTVHENVNKNEIK | 634-651 | 1 | N/D | N/D | N/D |
| LNSDSEAFLK | 761-770 | N/D | N/D | N/D | 1 |
| LNSDSEAFLKNPQR | 761-774 | N/D | N/D | N/D | 1 |
| FDQLFLEYK | 778-786 | 1 | N/D | N/D | 1 |
| QPNDIELDDTR | 837-847 | N/D | N/D | N/D | 1 |
| MLQNGINK | 882-889 | N/D | N/D | N/D | 1 |
| mNQNITLIQQIR | 905-916 | 1 | N/D | 1 | 1 |
| MLQSPLSAQNSR | 927-938 | 1 | 1 | 1 | 1 |
| mLQSPLSAQNSR | 927-938 | 1 | 1 | 1 | 1 |
| KIQPEESDSIVYK | 1159-1171 | 1 | N/D | N/D | N/D |
| VGAENDGDSSLFLR | 1285-1298 | 1 | 1 | N/D | N/D |

Peptides derived from Spt7p identified in the mass spectrometry analysis are summarized. The identified peptide sequences derived from Spt7 and their amino acid residue positions in Spt7 are shown in column 1 and 2, respectively. The numbers of peptide spectrum matches (PSMs) for each sample indicated in Table S1 are shown in columns 3 to 6. Peptide sequences are shown in the single letter code. The letters with lower case (‘c’ and ‘m’) indicate residues modified during the sample preparation for the mass spectrometry analysis. c: carbamidomethyl cysteine, m: oxidized methionine. N/D: not detected.

#### Table S3. Yeast strains used in this study

| Strain name | Genotype |
|---|---|
| BY4741 | *MATa his3Δ0 leu2Δ0 ura3Δ0 met15Δ0* |
| CPL33 | *MATa nup170Δ::KanMX* |
| TMY2837 | *MATa pep4Δ::HphMX* |
| TMY2838 | *MATa nup170Δ::KanMX pep4Δ::HphMX* |
| TMY2674-2B | *MATa PDR1-V5-HphMX HFI1-HA-HIS3MX* |
| TMY2674-14B | *MATa nup170Δ::KanMX PDR1-V5-HphMX HFI1-HA-HIS3MX* |
| TMY2716-6C | *MATa BAS2-V5-HphMX HFI1-HA-HIS3MX* |
| TMY2716-5B | *MATa nup170Δ::KanMX BAS2-V5-HphMX HFI1-HA-HIS3MX* |
| TMY2836 | *MATa SPT20-TAP-HIS3MX* |
| TMY2863 | *MATa nup170Δ::KanMX SPT20-TAP-HIS3MX* |
| TMY2854 | *MATa SPT7-TAP-HIS3MX* |
| TMY2821 | *MATa nup170Δ::KanMX SPT7-TAP-HIS3MX* |
| TMY2849 | *MATa pep4Δ::HphMX SPT7-TAP-HIS3MX* |
| TMY2846 | *MATa nup170Δ::KanMX pep4Δ::HphMX SPT7-TAP-HIS3MX* |
| TMY3168 | *MATa SPT20-TAP-HIS3MX SPT8-myc-KanMX RTG2-HA-HphMX* |
| TMY3170 | *MATa nup170Δ::NatMX SPT20-TAP-HIS3MX SPT8-V5-KanMX RTG2-HA-HphMX* |
| TMY3132 | *MATa his3::HIS3-*$P_{GAL1}$*-GFP-NLS-spt7(1088-1180)-GST-TCYC1* |
| TMY3134 | *MATa his3::HIS3-*$P_{GAL1}$*-GFP-NES-spt7(1088-1180)-GST-TCYC1* |
| TMY3136 | *MATa nup170Δ::KanMX his3::HIS3-*$P_{GAL1}$*-GFP-NLS-spt7(1088-1180)-GST-TCYC1* |
| TMY3138 | *MATa nup170Δ::KanMX his3::HIS3-*$P_{GAL1}$*-GFP-NES-spt7(1088-1180)-GST-TCYC1* |
| TMY3139 | *MATa pep4Δ::HphMX his3::HIS3-*$P_{GAL1}$*-GFP-NLS-spt7(1088-1180)-GST-TCYC1* |
| TMY3140 | *MATa pep4Δ::HphMX his3::HIS3-*$P_{GAL1}$*-GFP-NES-spt7(1088-1180)-GST-TCYC1* |
| TMY2995 | *MATa nup157Δ::URA3MX SPT20-TAP-HIS3MX* |
| TMY2996 | *MATa nup188Δ::URA3MX SPT20-TAP-HIS3MX* |
| TMY2997 | *MATa pom152Δ::URA3MX SPT20-TAP-HIS3MX* |
| TMY2862 | *MATa pep4Δ::HphMX SPT20-TAP-HIS3MX* |

#### Table S4. Oligo nucleotides used in this study

##### Oligos for RT-qPCR

| Oligo name | Sequence |
|---|---|
| TUB2F | TACTAGTGAAGGTATGGACGAATTG |
| TUB2R | TTCTTCATCATCTTCTACAGTAGCC |
| ACT1F | CATCCCATTTAACTGTAAGAAGAAT |
| ACT1R | GATCAGTCAATATAGGAGGTTATGG |
| SUR4-F | TGTTATGGTACTCAGGCTGC |
| SUR4-R | ACACCAGTAGAAGAACCGGA |
| ERG11-F | CAGATGATCTTGGCTGGACC |
| ERG11-R | CTTTCGGTGGTGGTAGACAC |
| DPS1-F | ATGGTCGTGGATACGTTGTG |
| DPS1-R | CGACAAATACGACACCGACT |
| DED81-F | TACCGATACCGTAACCACCA |
| DED81-R | GAATCGACGACATGGACGAA |
| RPL3-F | CTACCAGCTTCGACAGAACC |
| RPL3-R | GCTGACTTCTTCCAAAGCCT |
| PHO5-F | TCAAATGCACACCACGAGAA |
| PHO5-R | CATGTCCTGCTTGGGACTAC |
| HIS4-F | TCTAGACCCTCCTTCTTGGC |
| HIS4-R | TGCGGTGACTATTCAAGTGG |
| PDR5-F | TATGCGAATCATTTGGCGGA |
| PDR5-R | ACTTCAGCAATGGAGACACG |
| SNQ2-F | AAGTTGAAGTGTTGCGAGGT |
| SNQ2-R | TGAAGACGATGGATGCACTG |

##### Oligos for ChIP-qPCR

| Oligo name | Sequence |
|---|---|
| ACT1-2F | CAAGCGCTAGAACATACCAGA |
| ACT1-2R | TCCCCTTTCTACTCAAACCAA |
| ACT1-3F | ATGGAAGATGGAGCCAAAGC |
| ACT1-3R | CTGCCGGTATTGACCAAACT |
| RPL3-2F | TGTCTCTTTCGTGCTTCCTG |
| RPL3-2R | CTAAATGACCGTGACGTGGT |
| RPL3-3F | CTACCAGCTTCGACAGAACC |
| RPL3-3R | GCTGACTTCTTCCAAAGCCT |
| HIS4-1F | AGAATGCCCCCATCACAATC |
| HIS4-1R | GAGTCACTGTGCATGGGTTT |
| HIS4-2F | GCTCGAGCCATCCAAAAGTA |

| | |
|---|---|
| HIS4-2R | TTCACCTCCGATGTGTGTTG |
| HIS4-3F | TAGAACAAATCGGCAGCCTC |
| HIS4-3R | GGTTTGGTGGGGCTAGAATC |
| HIS4-4F | TCTAGACCCTCCTTCTTGGC |
| HIS4-4R | TGCGGTGACTATTCAAGTGG |
| PDR5-1F | GATCACGATTCAGCACCCTT |
| PDR5-1R | TACCACGGCGTAGAAGAGTT |
| PDR5-2F | TCTACGCCGTGGTACGATA |
| PDR5-2R | GTTTGCAACTTTGCGTGACT |
| PDR5-3F | TATGCGAATCATTTGGCGGA |
| PDR5-3R | ACTTCAGCAATGGAGACACG |
| PDR5-4F | TATGACTACCCCAAGTGCCA |
| PDR5-4R | CGTCTACGTTAGCAACACCA |
| SNQ2-1F | ACCTTCACGCCAGACTATGT |
| SNQ2-1R | ATGGGCGGACATTTAGTCAT |
| SNQ2-2F | ATTGTATTCCCTGCAACCCC |
| SNQ2-2R | GCATAAAAAGTGGTGAGGCG |
| ChrV-2F | TACTGACCTCCGAAGCTAGG |
| ChrV-2R | CAGGACTTTAGTCAGGACCG |
| ChrV-3F | TCTGCATTGTTCCCAACGAA |
| ChrV-3R | AAGGGACAGGCACTAAAACG |
| ChrII-1F | AATTACGGAAGCGCCAATGA |
| ChrII-1R | AACAAAAACGCGGAAGCTCT |

#### Supplemental Figure Legends

**Figure S1. The SAGA-dominated genes are overrepresented in genes showing the greatest downregulation in the *nup170Δ* strain.** A) Yeast protein encoding genes have been previously categorized based on their regulation by the transcriptional complexes TFIID and SAGA (Huisinga and Pugh, 2004). Plotted here are the percentage of genes in the indicated groups regulated primarily by the SAGA complex (SAGA-dominated, blue), TFIID (TFIID-dominated, green), or both complexes (SAGA/TFIID, purple), or were uncharacterized (no call, red). Three groups of genes are shown: all yeast protein-encoding genes (All) and the top 50 and 100 genes showing the greatest proportional decrease in mRNA levels in the *nup170Δ* strain relative to WT cells (based on data from Van De Vosse et al., 2013). B) The top 30 genes whose expression was most repressed in the *nup170Δ* mutant are listed, and their calculated mRNA ratios ([*nup170Δ*] / [WT]) are shown. Genes are colored in a same way as in (A) except for uncharacterized genes (no call), which are left uncolored.

### Figure S1

**A**

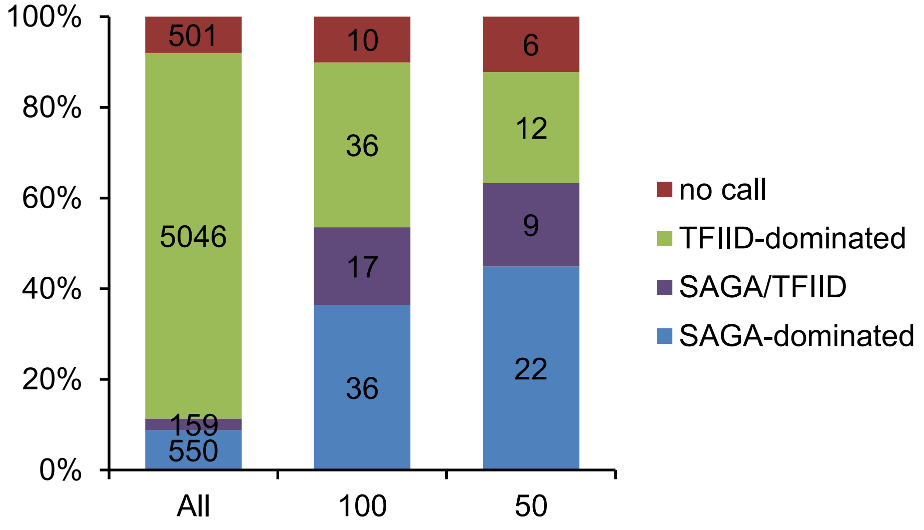

**B**

| Systematic name | Gene name | [*nup170Δ*]/[WT] | SAGA/TFIID |
|---|---|---|---|
| YBR093C | PHO5 | 0.1168 | SAGA/TFIID |
| YDR534C | FIT1 | 0.1767 | no call |
| YOL058W | ARG1 | 0.1820 | SAGA-dominated |
| YCR021C | HSP30 | 0.1975 | SAGA-dominated |
| YGL255W | ZRT1 | 0.2014 | SAGA/TFIID |
| YBR296C | PHO89 | 0.2268 | no call |
| YOR382W | FIT2 | 0.2368 | TFIID-dominated |
| YOR383C | FIT3 | 0.2479 | TFIID-dominated |
| YMR120C | ADE17 | 0.2684 | SAGA-dominated |
| YDR281C | PHM6 | 0.2756 | SAGA/TFIID |
| YLR346C | ORF:YLR346C | 0.2834 | SAGA-dominated |
| YAR071W | PHO11 | 0.2934 | TFIID-dominated |
| YNR060W | FRE4 | 0.2947 | SAGA-dominated |
| YLR302C | ORF:YLR302C | 0.2965 | TFIID-dominated |
| YHL047C | ARN2 | 0.2994 | TFIID-dominated |
| YJL052W | TDH1 | 0.3093 | SAGA-dominated |
| YJR150C | DAN1 | 0.3116 | SAGA/TFIID |
| YOL014W | ORF:YOL014W | 0.3171 | TFIID-dominated |
| YOR153W | PDR5 | 0.3202 | SAGA-dominated |
| YER011W | TIR1 | 0.3223 | SAGA/TFIID |
| YBR145W | ADH5 | 0.3232 | SAGA-dominated |
| YHR215W | PHO12 | 0.3287 | SAGA/TFIID |
| YMR006C | PLB2 | 0.3291 | SAGA/TFIID |
| YCL030C | HIS4 | 0.3315 | SAGA-dominated |
| YMR173W-A | ORF:YMR173W-A | 0.3334 | SAGA-dominated |
| YMR173W | DDR48 | 0.3498 | SAGA-dominated |
| YOR385W | ORF:YOR385W | 0.3530 | SAGA-dominated |
| YHR136C | SPL2 | 0.3666 | TFIID-dominated |
| YOL064C | MET22 | 0.3728 | TFIID-dominated |
| YOL016C | CMK2 | 0.3735 | SAGA-dominated |
